# Rapid ecosystem change after the cod collapse in Newfoundland & Labrador

**DOI:** 10.1101/2024.10.22.619726

**Authors:** Alannah Wudrick, Paul M. Regular, Jamie C. Tam, David Bélanger, Tyler D. Eddy

## Abstract

The cod collapse of Newfoundland and Labrador in the early 1990s was predicated by collapses of the key forage fish species in the region, capelin, and the wider groundfish community. After these collapses, rapid changes in ecosystem structure and function have been observed. From the early 2000s to mid 2010s, invertebrates dominated ecosystem biomass and the snow crab fishery became important to the region. During the late 2010s, invertebrates declined as groundfish stocks increased, although not to pre collapse levels. We developed two ecosystem models representing 1) the Grand Banks and 2) the Newfoundland and Labrador Shelf ecosystem structure and function for 2018-2020. We compared these models with existing ecosystem models that represent the combined Grand Banks and Newfoundland and Labrador Shelf ecosystem for the pre collapse (1985-1987) and invertebrate dominated (2013-2015) periods. Variability in primary productivity, species biomass, and catch reveal that the two model regions are distinct in ecosystem structure and function. Primary and secondary production has increased in the most recent period, however capelin biomass – an important factor for cod recovery – has remained low. Harp seals play a strong top-down ecosystem role and may be contributing to impaired cod recovery. Since the decline in capelin abundance, Arctic cod became an important alternative prey on the Newfoundland and Labrador Shelf region compared to reliance on sand lance and squid on the Grand Banks. Capelin has not shown signs of recovery in either region. Overall, the ecosystem structure in 2018-2020 in both regions is closer to the pre-collapse state than it was in 2013-2015, and the rapid change indicates that the system is highly dynamic.

## 1. Introduction

Marine ecosystems are affected by bottom-up processes from environmental conditions and top-down forces from predators and fishing – both of which influence ecosystem structure and function (1). The combined effects of fisheries and climate change can cause rapid changes in marine ecosystem structure and function on large scales (2,3). Highly dynamic ecosystems can be difficult to predict, but management approaches that incorporate climate and environmental variables can help detect and anticipate changes in ecosystem structure and function (4,5). Ecosystem-based fisheries management (EBFM) seeks to balance multi-species fishery yield, socioeconomic wellbeing, and ecosystem sustainability under a comprehensive ecosystem management framework, as opposed to multiple independent single-species stock assessments (6,7). The ecosystem approach to fisheries management (EAFM) applies ecosystem considerations within single species stock assessments (8,9). EAFM and EBFM are better suited than traditional approaches to fisheries management to respond to climate events such as marine heat waves and other climate change impacts on dynamic marine ecosystems (10).

The Newfoundland and Labrador marine ecosystem, a dynamic system, experienced dramatic shifts in structure and function during the early 1990s. The collapse of key species such as capelin (*Mallotus villosus*) and Atlantic cod (*Gadus morhua*), along with other commercial and non-commercial fish species, triggered significant ecological reorganization (11). The major factors that contributed to the groundfish collapse were overfishing in combinations with unusually low sea surface and bottom temperatures (3). In response to this collapse, a groundfish fishing moratorium was implemented in 1992, prohibiting the catch of Atlantic cod and other species in the region including American plaice (*Hippoglossoides platessoides*), haddock (*Melanogrammus aeglefinus*), and redfish (*Sebastes fasciatus*). Subsequently, invertebrate populations such as snow crab (*Chionoecetes opilio*) and Northern shrimp (*Pandaulus borealis*) boomed between the mid 1990’s and early 2000’s, and many fishers transitioned to fishing invertebrates.

There were several hypotheses as to why the Atlantic cod stock was struggling to recover. It was suggested that the growing harp seal (*Pagophilus groenlandicus*) population hindered cod recovery through high levels of consumption (12). Capelin are a key driver of cod population dynamics (13), meaning the reduced availability of forage species like capelin was thought to negatively impact cod (11). Capelin population dynamics are not fully understood but have been linked to the timing and intensity of the spring phytoplankton bloom, which influences the abundance of nutrient-rich zooplankton that capelin consume and provide energy to higher trophic levels (14). Though capelin continue to exhibit low productivity (15), modest growth has supported a partial recovery of cod (13,16), leading to the end of the commercial fishing moratorium for the most iconic Atlantic cod stock: Northern cod (17). In parallel, the dominance of the invertebrate community has dwindled through the late 2010’s to the early 2020’s, and the rest of the groundfish community has shown a modest recovery in a relatively short period of time.

Highly dynamic ecosystems pose challenges for fisheries management, as fished species biomasses can change rapidly in response to the environment (5). Improving our understanding of ecosystem structure and function can benefit single species stock assessments in addition to EAFM and EBFM goals, as fisheries management can better plan or prepare for dynamic periods (18). Evaluating ecosystem structure and function requires ecosystem scale tools such as ecosystem models (19). Ecopath ecosystem models have been developed for the Newfoundland and Labrador region, representing the pre-collapse ecosystem in 1985-1987 (20) and the high invertebrate biomass ecosystem in 2013-2015 (21). These models considered the Newfoundland and Labrador Shelf and Grand Banks as one model domain, despite these regions having distinct differences in bathymetry, productivity, biodiversity, and climate (22). What remains unknown is how ecosystem structure and function vary 1) between these two regions and 2) over time in recent years as groundfish populations have increased and invertebrates have declined. This study develops two new ecosystem models to represent the ecosystem structure and function of the Newfoundland and Labrador Shelf and Grand Banks separately, for 2018-2020. We investigate how ecosystem structure and function, abundance of key species, and fisheries catches vary in space and time.

## 2. Methods

### 2.1 Study area

The Newfoundland and Labrador Shelf, in the Northwest Atlantic Ocean of Canada, is a stretch of underwater rocky continental shelves that extend offshore to approximately 200 nautical miles (nm; 370 km), averaging 200-500 m depth (23) (Fig 1). Oceanographically, the shelves are influenced by cold-water flow from the Labrador current, which transports freshwater glacier melt from the Arctic southward along the coast. This mass of cold water (known at the cold intermediate layer, or CIL) is a strong driver of environmental conditions on the shelf, and varies both seasonally and interannually (24). The dynamic vertical stratification of the CIL strongly influences primary productivity and the horizontal distribution of zooplankton in the area, with different species assemblages found in different water masses across the shelf (25).

**Fig 1.**
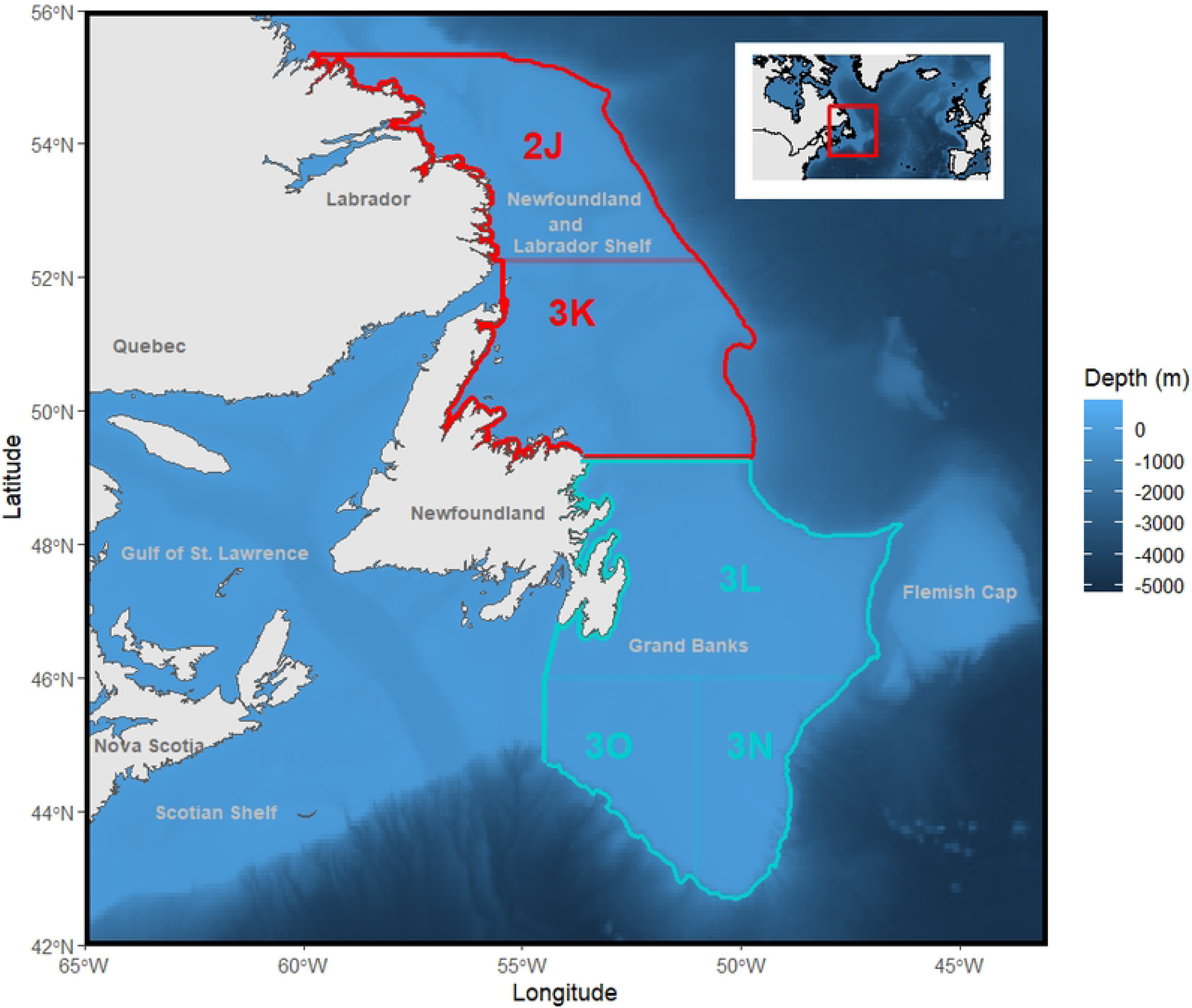
Map of the model domains. Grand Banks (light blue; Northwest Atlantic Fisheries Organization (NAFO) divisions 3LNO) and the Newfoundland and Labrador Shelf (red; NAFO divisions 2J3K). Model domains extend from shore to the 1000 m isobath.

The Grand Banks are a series of soft, sandy and muddy banks that average 50-100 m in depth and are used for fishing and oil and gas production (26) (Fig 1). Flow from the Labrador Current reaches the north of the Grand Banks, where it is met by a southern warm flow of water from the Gulf Stream, which influences nutrient mixing and promotes primary and secondary productivity for the region (27). The ecosystem models developed for 1985-1987 (20) and 2013-2015 (21) represented the Northwest Atlantic Fisheries Organization (NAFO) Divisions 2J3KLNO to 1000 m depth (Fig 1). The ecosystem models developed in this study represent the Grand Banks (3LNO) and Newfoundland and Labrador Shelf (2J3K) separately, also to 1000 m depth, for 2018-2020. These areas align with many fisheries stock boundaries and Canada’s Exclusive Economic Zone (EEZ) that extends out to 200 nautical miles (Fig 1). The area of the 2J3KLNO model domain to the 1000 m isobath is 495 000 km^2^, the Newfoundland and Labrador Shelf (2J3K) is 237 600 km^2^, and the Grand Banks (3LNO) is 257 400 km^2^.

### 2.2 Ecosystem models

The Newfoundland and Labrador Shelf (2J3K) and Grand Banks (3LNO) ecosystems were modeled using Ecopath with Ecosim (EwE) version 6.6.8 software (28). EwE employs a mass-balance approach to track energy and biomass flow and represent instantaneous snapshots of ecosystem structure and function. The mass balance approach requires sufficient biomass at the bottom of the food web to sustain biomass at the top of the food web based on production and consumption rates of functional groups. The core EwE routine is based on a series of linear equations that solve for one unknown. Along with diet information, three of four parameters must be known: *B* (biomass), *P/B* (production), *Q/B* (consumption), and *EE* (ecotrophic efficiency; proportion of biomass of prey consumed). Species-specific production is calculated as:

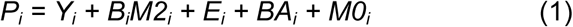

Where *P_i_* = production rate of species *i*; *Y_i_* = total fishery catch rate of species *i; B_i_* = total biomass of species *i; M2_i_* = predation mortality on species *i; E_i_* = net migration (emigration - immigration) of species *i; BA_i_* = biomass accumulation rate for species *i; M0_i_* = all other mortality not accounted for by predation or catch of species *i*. All rates are averaged annually. Biomass accumulation (BA) and net migration (E) describe the change in biomass over time and the emigration rate minus the immigration rate respectively. However, these terms were not included in either model. (i.e., any accumulation of biomass was assumed to be consumed and net migration was assumed to be 0). Therefore, in order for the models to be in mass balance, the following condition must be met based on equation 1:

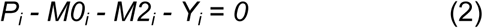

*M0i* can be expressed as (1-*EEi*), or, the proportion of biomass not consumed by higher trophic levels. *M2i* is calculated by summing the annual consumption of species *i* by all *j* predator groups:

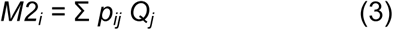

Where: *p_ij_* = Proportion by mass of predators *j*’s diet that is comprised of prey *i*; *Q_j_* = Annual consumption of biomass by predator *j*. Annual consumption (*Q_i_*) can be calculated by: Consumption = production + respiration + unassimilated food, or:

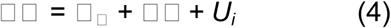

Where: *P_i_*= Production rate of species *i*; *R_i_* = Respiration rate of species *i*; *U_i_* = Proportion of food not assimilated into body tissue.

To reduce model complexity, species of similar functional type were combined into functional groups following previous ecosystem models (20,21). Functional groups can be age-structured, referred to as multi-stanza groups in EwE. Multi-stanza groups have two main assumptions: 1) the growth of a species follows a von Bertalanffy growth curve (weight is proportional to length cubed) and 2) the population has a relatively stable mortality and recruitment rate. Multi-stanza groups are useful in situations where predation mortality and/or catch differs significantly between juveniles and adults of the same species (28). In both models, Atlantic cod and American plaice are multi-stanza groups.

### 2.3 Data sources and model balancing

Biomass (*B*) estimates were obtained from Fisheries and Oceans Canada (NAFO) and NAFO stock assessments and research vessel (RV) stratified random sampling surveys (S1 and S2 Table). Survey data are analyzed using a standard stratified analysis via a local R package called *Rstrap* (For details on methodology, see (29)). Production to biomass (*P/*B) and consumption to biomass (*Q/*B) ratios were estimated from the literature (S1 and S2 Table). In areas where consumption rates were not available for species in the model area, an assumption of a production to consumption ratio, *P/Q* (also known as ‘growth efficiency’) of 0.15 was used, following best practice (30). Diet composition was estimated using RV diet survey data and relevant literature (Table S3 and S4). Catch data for commercial finfish and shellfish fisheries were obtained from the NAFO STATLANT 21A database (NAFO 2016). The pedigree function in Ecopath is a user-defined score that indicates parameter data quality from high-precision local sampling (low uncertainty), to ‘guestimates’ (high uncertainty) (S5 and S6 Table). Generally, models were balanced by modifying parameters with low data pedigree and high uncertainty first, such as diet data, as opposed to parameters with low uncertainty such as biomass estimates (initial and balanced model parameters shown in Tables 1 and 2, S7 and S8 Table respectively).

**Table 1.** Balanced 2018-2020 Newfoundland and Labrador Shelf model with biomass (*B*), production (*P/B*), consumption (*Q/B*), ecotrophic efficiency (*EE*), catch, fishing mortality (*F*) and natural mortality (*M*) parameter estimates. Bolded values were estimated by Ecopath. Shaded rows indicate size structured groups.

| Functional group | Biomass<br>(t/km<sup>2</sup>) | *P/B* | *Q/B* | *EE* | Trophic<br>level | Catch<br>(t) | *F* | *M* |
| --- | --- | --- | --- | --- | --- | --- | --- | --- |
| Whale fish eater | 0.700 | 0.12 | 6.00 | **0.00** | **4.30** | 0.00 | **0.00** | **3.54E-4** |
| Whale zooplankton eater | 0.014 | 0.06 | 3.47 | **0.50** | **3.59** | 0.00 | **0.00** | **0.027** |
| Whale squid eater | 0.121 | 0.09 | 6.00 | **0.00** | **4.53** | 0.00 | **0.00** | **0.00** |
| Whale mammal eater | 3.06E-4 | 0.08 | 8.10 | **0.00** | **5.25** | 0.00 | **0.00** | **0.00** |
| Seal harp | 0.380 | 0.15 | 17.64 | **0.59** | **4.49** | 0.008 | **0.031** | **0.005** |
| Seal hooded | 0.032 | 0.12 | 18.33 | **0.14** | **4.86** | 0.00 | **0.00** | **0.016** |
| Seal other | 0.016 | 0.13 | 13.00 | **0.03** | **4.69** | 0.00 | **0.00** | **0.004** |
| Seabird piscivore | 0.007 | 0.25 | 119.0 | **0.07** | **4.28** | 0.00 | **0.00** | **0.018** |
| Seabird planktivore | 0.006 | 0.15 | 64.61 | **0.00** | **3.48** | 0.00 | **0.00** | **0.00** |
| Seabird benthivore | 0.002 | 0.13 | 45.29 | **0.00** | **3.63** | 0.00 | **0.00** | **0.00** |
| Greenland shark | 0.009 | 0.01 | 0.13 | **0.00** | **5.18** | 0.00 | **0.00** | **0.00** |
| Atlantic cod >35cm | 2.180 | 0.40 | 1.62 | **0.97** | **4.05** | 0.018 | **0.008** | **0.390** |
| Atlantic cod <=35cm | **0.200** | 0.40 | **4.10** | **0.99** | **3.67** | 0.003 | **0.016** | **0.381** |
| Greenland halibut | 0.800 | 0.64 | 2.40 | **0.87** | **4.42** | 0.007 | **0.009** | **0.541** |
| Other piscivorous fish | 0.037 | 0.31 | 4.95 | **0.94** | **4.38** | 6.45E-5 | **0.002** | **0.289** |
| Redfish | 1.20 | 0.33 | 2.00 | **0.89** | **3.71** | 1.30E-4 | **1.08E-4** | **0.264** |
| Arctic cod | 4.548 | 0.34 | 4.00 | **0.99** | **3.48** | 0.00 | **0.00** | **0.331** |
| Other plank-piscivorous fish | 0.028 | 0.35 | 2.30 | **0.79** | **3.56** | 0.00 | **0.00** | **0.276** |
| American plaice >35cm | 0.40 | 0.60 | 2.00 | **0.73** | **3.65** | 1.54E-5 | **3.85E-5** | **0.330** |
| American plaice <= 35cm | **1.411** | 0.60 | **3.86** | **0.99** | **3.59** | 0.00 | **0.00** | **0.525** |
| Thorny skate | 0.130 | 0.15 | 1.79 | **1.00** | **3.98** | 1.80E-5 | **1.38E-4** | **0.149** |
| Other large benthivorous fish | 0.985 | 0.30 | 1.33 | **0.83** | **3.63** | 6.31E-6 | **7.01E-6** | **0.261** |
| Witch flounder | 0.060 | 0.38 | 2.60 | **0.83** | **3.49** | 2.60E-4 | **0.004** | **0.309** |
| Other medium benthivorous fish | 0.385 | 0.40 | 2.00 | **0.96** | **3.66** | 0.00 | **0.00** | **0.376** |
| Small benthivorous fish | **2.543** | 0.45 | 2.00 | 0.98 | **3.60** | 0.00 | **0.00** | **0.441** |
| Herring | **2.211** | 1.15 | 3.15 | 0.98 | **3.67** | 0.01 | **0.005** | **1.12** |
| Sandlance | **1.935** | 1.15 | 7.66 | 0.98 | **3.56** | 0.00 | **0.00** | **1.13** |
| Capelin | 3.400 | 1.15 | 4.30 | **0.93** | **3.48** | 0.011 | **0.004** | **1.13** |
| Other planktivorous fish | **3.061** | 1.15 | 4.19 | 0.98 | **2.89** | 0.02 | **0.007** | **1.12** |
| Squid | 0.400 | 3.40 | 13.20 | **1.00** | **3.56** | 0.005 | **0.014** | **3.38** |
| Shrimp | 0.585 | 1.70 | 11.33 | **0.99** | **2.44** | 0.047 | **0.080** | **1.59** |
| Snow crab | 0.242 | 0.50 | 3.06 | **0.95** | **3.56** | 0.013 | **0.053** | **0.403** |
| Predatory invertebrates | 20.0 | 2.70 | 8.73 | **0.78** | **3.16** | 0.002 | **8.81E-5** | **2.08** |
| Deposit feeding invertebrates | 85.0 | 3.00 | 9.10 | **0.77** | **2.46** | 1.00E-4 | **1.18E-6** | **2.31** |
| Suspension feeding invertebrates | 61.0 | 1.50 | 3.70 | **0.56** | **2.00** | 3.09E-5 | **5.07E-7** | **0.831** |
| Macrozooplankton | **17.88** | 3.43 | 19.50 | 0.98 | **2.48** | 0.00 | **0.00** | **3.36** |
| Large mesozooplankton | 14.0 | 10.0 | 28.00 | **0.87** | **2.58** | 0.00 | **0.00** | **8.59** |
| Small mesozooplankton | 5.50 | 33.0 | 105.4 | **0.88** | **2.37** | 0.00 | **0.00** | **28.8** |
| Microzooplankton | 5.360 | 73.0 | 240.0 | **0.93** | **2.00** | 0.00 | **0.00** | **67.8** |
| Large phytoplankton | 7.175 | 107.0 |  | **0.88** | **1.00** | 0.00 | **0.00** | **94.1** |
| Small phytoplankton | 11.79 | 107.0 |  | **0.93** | **1.00** | 0.00 | **0.00** | **99.4** |
| Detritus | 1.00 |  |  | **0.91** | **1.00** | 0.00 | **0.00** | **0.00** |

**Table 2.** Balanced 2018-2020 Grand Banks model with biomass (*B*), production (*P/B*), consumption (*Q/B*), ecotrophic efficiency (*EE*), catch, fishing mortality (*F*) and natural mortality (*M*) parameter estimates. Bolded values were estimated by Ecopath. Shaded rows indicate size structured groups.

| Functional group | Biomass<br>(t/km<sup>2</sup>) | *P/B* | *Q/B* | *EE* | Trophic<br>level | Catch<br>(t) | *F* | *M* |
| --- | --- | --- | --- | --- | --- | --- | --- | --- |
| Whale fish eater | 0.730 | 0.12 | 6.00 | **0.00** | **4.35** | 0.00 | **0.00** | **3.13E-4** |
| Whale zooplankton eater | 0.016 | 0.06 | 3.47 | **0.38** | **3.58** | 0.00 | **0.00** | **0.021** |
| Whale squid eater | 0.112 | 0.09 | 6.00 | **0.00** | **4.58** | 0.00 | **0.00** | **0.00** |
| Whale mammal eater | 2.82E-4 | 0.08 | 8.10 | **0.00** | **5.29** | 0.00 | **0.00** | **0.00** |
| Seal harp | 0.380 | 0.15 | 17.64 | **0.53** | **4.50** | 0.007 | **0.029** | **0.005** |
| Seal hooded | 0.007 | 0.12 | 18.33 | **0.56** | **5.01** | 0.00 | **0.00** | **0.064** |
| Seal other | 0.014 | 0.13 | 13.00 | **0.03** | **4.78** | 0.00 | **0.00** | **0.005** |
| Seabird piscivore | 0.007 | 0.25 | 119.4 | **0.06** | **4.36** | 0.00 | **0.00** | **0.016** |
| Seabird planktivore | 0.006 | 0.15 | 64.60 | **0.00** | **3.52** | 0.00 | **0.00** | **0.00** |
| Seabird benthivore | 0.002 | 0.13 | 45.29 | **0.00** | **3.64** | 0.00 | **0.00** | **0.00** |
| Greenland shark | 0.009 | 0.01 | 0.13 | **0.00** | **5.27** | 0.00 | **0.00** | **0.00** |
| Atlantic cod >35cm | 0.568 | 0.40 | 1.17 | **0.48** | **4.45** | 0.018 | **0.032** | **0.350** |
| Atlantic cod <=35cm | **0.052** | 0.40 | **2.96** | **0.69** | **3.69** | 0.003 | **0.064** | **0.311** |
| Greenland halibut | 0.200 | 0.64 | 2.30 | **0.53** | **4.44** | 0.002 | **0.013** | **0.352** |
| Silver hake / Pollock | 0.128 | 0.40 | 4.10 | **0.65** | **4.32** | 0.000 | **0.003** | **0.312** |
| Other piscivorous fish | 0.122 | 0.38 | 1.65 | **0.99** | **4.48** | 0.008 | **0.066** | **0.305** |
| Redfish | 1.248 | 0.33 | 2.00 | **0.88** | **3.74** | 0.073 | **0.058** | **0.194** |
| Arctic cod | 1.050 | 0.60 | 4.00 | **0.95** | **3.52** | 0.00 | **0.00** | **0.405** |
| Other plank-piscivorous fish | 0.028 | 0.35 | 2.30 | **0.64** | **3.63** | 7.77E-6 | **2.81E-4** | **0.225** |
| American plaice >35cm | 0.830 | 0.50 | 2.00 | **0.62** | **3.91** | 0.004 | **0.005** | **0.225** |
| American plaice <= 35cm | **1.783** | 0.51 | **3.73** | **0.56** | **3.83** | 6.00E-5 | **3.36E-5** | **0.221** |
| Thorny Skate | 0.709 | 0.20 | 1.79 | **0.45** | **4.31** | 0.011 | **0.016** | **0.061** |
| Haddock | 0.039 | 0.21 | 2.08 | **0.96** | **3.75** | 2.00E-4 | **0.005** | **0.199** |
| Other large benthivorous fish | 0.655 | 0.30 | 1.33 | **0.52** | **3.66** | 0.001 | **0.002** | **0.253** |
| Yellowtail flounder | 1.180 | 0.39 | 3.60 | **0.52** | **3.97** | 0.017 | **0.015** | **0.188** |
| Witch flounder | 0.063 | 0.20 | 2.60 | **0.92** | **3.50** | 0.003 | **0.048** | **0.133** |
| Other medium benthivorous fish | 0.320 | 0.40 | 2.00 | **0.74** | **3.67** | 1.58E-4 | **4.94E-4** | **0.339** |
| Small benthivorous fish | 1.618 | 0.45 | 2.00 | 0.99 | **3.62** | 6.47E-6 | **4.09E-6** | **0.445** |
| Herring | 1.893 | 1.15 | 3.15 | 0.98 | **3.69** | 0.003 | **0.002** | **1.13** |
| Sandlance | 4.529 | 1.15 | 7.66 | 0.98 | **3.58** | 0.00 | **0.00** | **1.13** |
| Capelin | 3.770 | 1.20 | 4.30 | **0.91** | **3.51** | 0.019 | **0.006** | **1.17** |
| Other planktivorous fish | **2.886** | 1.15 | 4.19 | 0.98 | **2.90** | 1.79E-4 | **6.38E-5** | **1.13** |
| Squid | 0.700 | 3.40 | 13.20 | **0.78** | **3.55** | 0.005 | **0.009** | **2.58** |
| Shrimp | 0.666 | 1.70 | 11.30 | **0.97** | **2.45** | 0.00 | **0.00** | **1.60** |
| Snow crab | 0.430 | 0.50 | 3.06 | **0.99** | **3.58** | 0.020 | **0.046** | **0.450** |
| Predatory invertebrates | 20.0 | 2.70 | 8.73 | **0.87** | **3.16** | 0.001 | **6.65E-5** | **2.31** |
| Deposit feeding invertebrates | 85.0 | 3.00 | 10.00 | **0.83** | **2.48** | 4.00E-4 | **4.71E-6** | **2.49** |
| Suspension feeding invertebrates | 61.0 | 1.50 | 3.70 | **0.93** | **2.00** | 0.064 | **0.001** | **1.39** |
| Macrozooplankton | 21.0 | 3.20 | 19.5 | 0.98 | **2.50** | 0.00 | **0.00** | **3.26** |
| Large mesozooplankton | 22.9 | 8.20 | 28.0 | **0.99** | **2.59** | 0.00 | **0.00** | **8.92** |
| Small mesozooplankton | 7.816 | 31.0 | 105.4 | **0.92** | **2.45** | 0.00 | **0.00** | **31.6** |
| Microzooplankton | 7.470 | 72.0 | 240 | **0.94** | **2.00** | 0.00 | **0.00** | **69.3** |
| Large phytoplankton | 8.780 | 106 |  | **0.96** | **1.00** | 0.00 | **0.00** | **74.6** |
| Small phytoplankton | 15.800 | 106 |  | **0.97** | **1.00** | 0.00 | **0.00** | **75.4** |
| Detritus | 1.000 |  |  | **0.93** | **1.00** |  |  | **3.13E-4** |

### 2.4 Functional groups

The Grand Banks model had 45 defined functional groups and the Newfoundland and Labrador Shelf model had 42 (S9 Table). Functional groups represented fish (19 groups for the Newfoundland and Labrador Shelf; 22 groups for the Grand Banks), cetaceans (4 groups), seals (3 groups), seabirds (3 groups), invertebrates (6 groups), and plankton (6 groups). Where applicable, the same functional groups from the previous models were used to facilitate model comparison ((20,21)). The microbial loop functional groups were removed due to data limitations. The minke whale group had originally been defined for comparisons with the Barents Sea ecosystem (21) but was combined with the fish-eating whales functional group for the 2018-2020 models.

#### 2.4.1 Cetaceans

Cetaceans were represented by four functional groups based on primary prey type: fish eaters, zooplankton eaters, squid eaters, and mammal eaters. Some whale species are seasonally transient while others stay within the model regions year-round (31–33). Cetacean biomass was calculated by multiplying estimated population size by estimated average body weight scaled by residency time where applicable and divided by the model area. Residency times for migratory species were 180 days/year to incorporate the spring and summer seasons when sightings occur most frequently (34). Residency time for non-migratory species was estimated as 270 days to account for seasonal movements on and off the shelf (34,35).

Population estimates were obtained from the Northwest Atlantic International Sightings Survey (NAISS) marine mammal census survey, Committee on the Status of Endangered Wildlife in Canada (COSEWIC) assessments, National Oceanic and Atmospheric Administration (NOAA) stock assessment reports (36–41). Mean body weights for cetaceans were estimated from the literature (32,42–45). *P/B* ratios for cetaceans were estimated using Hoenig’s equation for marine mammal mortality (46), based on maximum age (45). *Q/B* ratios were calculated from annual consumption estimates found in the literature (32,45,47). No commercial catch of cetaceans occurs within the study area. Some species such as narwal and beluga are hunted by Indigenous groups for food, social, and ceremonial purposes, but this occurs outside of the study area.

#### 2.4.2 Seals

Seals were represented by three seal functional groups: harp seals (*Pagophilus groenlandicus*), hooded seals (*Cystophora cristata*), and other seals which contained grey seals (*Halichoerus grypus*) and harbour seals (*Phoca vitulina*). Biomass estimates for harp seals were obtained from DFO stock assessments (48) and scaled using a residency time of 182 days to represent their seasonal migrations in and out of the study area (49,50). Biomass estimates for hooded seals were based on tagging and migration studies, hooded seal biomass was scaled using a residency time of 180 days for both areas, with 80% of biomass on the Newfoundland and Labrador Shelf and 20% on the Grand Banks (51,52). *P/B* ratios for seals were calculated using Hoenig’s equation for marine mammal mortality (46), based on maximum age (45). Seal consumption was estimated from seal consumption models (21). Only harp seals had reported catches in the study area and were not reported by location, so catches were evenly split between the Newfoundland and Labrador Shelf and the Grand Banks (48).

#### 2.4.3 Seabirds

Seabirds were grouped based on diet; planktivore, piscivore, and benthivore. Seabird abundance was estimated from annual surveys conducted by Canadian Wildlife Services for the Seabirds at Sea program (53). The abundance of seabirds within the study area varies throughout the year. Seasonally, large numbers of birds move in and out of the study. To account for these movements, biomass was scaled by residency time. *P/B* for seabird groups was assumed to be 0.25 (20). *Q/B* for piscivorous seabirds was based on a consumption study spanning NAFO subareas 2 and 3 (54). Planktivorous seabird *Q/B* was based on a consumption study of dovekies (*Alle alle*) (55). Benthivorous seabird *Q/B* was estimated from the consumption of common eiders (*Somateria mollissima*) in the Gulf of St. Lawrence (56).

#### 2.5.4 Fish

Most single species fish functional group biomasses were estimated from DFO stock assessment documents (S1 and S2 Tables). Biomass for fish functional groups that represented multiple species was estimated from RV survey data. For forage fish such as small benthivorous fish, herring, sandlance, and other planktivorous fish, Ecopath estimated biomass using an *EE* of 0.98 following best practice (19,21). *P/B* was derived from estimates of natural mortality using Hoenig’s equation for fish mortality (46). *Q/B* was estimated from local consumption studies where possible, otherwise *P/Q* was 0.15 yr^-1^ (S1 and S2 Tables). Commercial catch data for all groups were obtained from the NAFO STATLANT 21A database to estimate catches (57). Diets were informed by RV survey data and other localized diet studies where applicable (S1 and S2 tables).

#### 2.4.5 Invertebrates

Biomass and *P/B* estimates for snow crab and Northern shrimp were obtained from DFO and NAFO stock assessments (58–60). *Q/B* was estimated based on a growth efficiency of 0.15 (20,21). Biomass for the squid group, which contained all cephalopods but was predominately Northern shortfin squid (*Illex illecebrosus*), was estimated using RV survey data. *P/B* for squid estimated at 0.07 week^-1^ (61) and *Q/B* was estimated using consumption values of squid from the Northeastern United States (62). All other invertebrates were grouped based on feeding strategy (S9 Table). Predatory invertebrates include lobster, other crab species, and other crustaceans (S9 Table). Deposit feeding invertebrates include urchins, sand dollars, polychaetes, chaetognaths, and isopods, and the suspension feeding invertebrates functional group includes sponges, corals, bivalve molluscs, sea anemones, brittle / basket stars, and ascidians (S5 Table). There was limited target survey data for these aggregated groups, so biomass estimates for all three groups were based on RV survey data and grab sample surveys from 2008-2010 (63). *P/B* ratios for predatory invertebrates were based on estimates from a Scotian Shelf model (64). The *P/B* ratios for deposit feeding invertebrates and suspension feeding invertebrates were averaged from production estimates of molluscs and echinoderms (21,65–68). *Q/B* ratios were based on the assumption that the *P/Q* ratio is 0.15 yr^-1^. Commercial catch data for all groups was estimated from the NAFO STATLANT 21A database (57).

#### 2.4.6 Zooplankton

Macrozooplankton biomass was estimated by Ecopath using an *EE* of 0.98 (21). For all other zooplankton groups, abundance was estimated using data from the Atlantic zone monitoring program (69). The dry weight of the zooplankton species at different life stages was estimated from a literature review and was converted to wet weight using a conversion factor (70). The resulting estimates were compared to the 2013-2015 model and adjusted with the understanding that zooplankton biomass increased since 2015 in both areas (24). Microzooplankton *P/B* was estimated using an energy model for the Northeastern United States (71). Small mesozooplankton *P/B* was estimated from a daily production rate empirical equation (72). Large mesozooplankton *P/B* was estimated based on the production of *Calanus finmarchicus* on the Scotian Shelf (73), and macrozooplankton *P/B* was estimated from euphausiid production studies (20,74). Macrozooplankton *Q/B* was based on the consumption of three euphausiid species in the Gulf of St. Lawrence (75). Large mesozooplankton, small mesozooplankton, and microzooplankton *Q/B* was estimated using a P/Q of 0.3 yr^-1^ (30).

#### 2.4.7 Phytoplankton

The phytoplankton functional groups are size structured. Large phytoplankton includes all microphytoplankton (∼20-200 µm) and small phytoplankton includes nano and pico phytoplankton (∼0.2-20 µm). Biomass estimates were derived from remote sensing data using PhytoFit R shiny application (76). Surface layer mean chlorophyll α concentrations were obtained from Visible Infrared Imaging Radiometer Suite (VIIRDS) satellite images with 4 km resolution (method Poly4) and separated by cell size, to give estimates of weekly large and small phytoplankton derived chlorophyll α for 2018-2020. Chlorophyll α concentrations were converted to biomass estimates using a monthly chlorophyll: carbon conversion ratio (77).

Phytoplankton *P/B* was the same as the 2013-2015 model, which used primary production estimates for the Newfoundland Shelf (provided by Carla Caverhill, unpublished data).

### 2.5 Model analyses

Summary statistics such as net primary production (NPP), net system production, total system throughput (TST), omnivory index, and connectance index were calculated by Ecopath. A Lindeman spine was used to examine trophic flows and pathways (78). The mixed trophic impact (MTI) analysis indicates how changes in biomass of one functional group will affect other functional groups (19). Estimates of fishing mortality (*F*) and predation mortality (*M*) were estimated by Ecopath based on catch and consumption data.

## 3. Results

### 3.1 Model balancing

Initial parameter estimates (S7 and S8 Table) did not produce a balanced model and adjustments were made to achieve mass balance. Adjusting diet matrices was the main strategy to balance both models. The largest changes in diet were made to witch flounder (*Glyptocephalus cynoglossus*), decreasing their consumption of predatory invertebrates and increasing their consumption of deposit feeding invertebrates (S3 and S4 Fig). The largest parameter changes to both models were made to increase the biomass of squid, other large benthivorous fish, and other medium benthivorous fish (S1 and S2 Figs). Zooplankton biomass was increased for the Grand Banks, but not for the Newfoundland and Labrador Shelf. The largest decreases in biomass were for small benthivorous fish in both regions. Small benthivorous fish biomass was estimated by Ecopath, which produced an unrealistically high estimate of 43 t/km^2^ due to predation from other large groups, primarily predatory invertebrates (S7 and S8 Table). When predation levels were adjusted, the estimate fell by 1500% to 2.8 and 1.8 t/km^2^ on the Newfoundland and Labrador Shelf and Grand Banks, respectively (S1 and S2 Figs).

### 3.2 Spatial comparisons

#### 3.2.1 Ecosystem structure and function

Functional groups ranged from trophic level 1 (primary producers) to 5.25 (whale mammal eater) on the Newfoundland and Labrador Shelf and 5.30 (whale mammal eater) on the Grand Banks (Tables 1 and 2). Both models had the same six functional groups with the highest trophic levels; mammal eating whales, Greenland shark, hooded seals, other seals, squid eating whales, and harp seals, respectively. Functional group trophic levels were always higher for the Grand Banks than the Newfoundland and Labrador Shelf, with the largest difference observed for Atlantic cod >35 cm, which had a trophic level of 4.05 on the Newfoundland and Labrador Shelf and 4.45 on the Grand Banks. The next largest differences were thorny skate (3.98 Newfoundland and Labrador Shelf and 4.31 Grand Banks), both American plaice size groups (3.65 Newfoundland and Labrador Shelf and 3.90 Grand Banks >35 cm; 3.59 Newfoundland and Labrador Shelf and 3.84 Grand Banks), and hooded seals (4.86 Newfoundland and Labrador Shelf and 5.01 Grand Banks). The rest of the trophic differences were under 0.1, with the majority (70%) less than 0.05.

In both regions, most of the energy flow occurred between the first three trophic levels (82.8% in Newfoundland and Labrador Shelf and 77.2% in Grand Banks) (S5 and S6 Fig). Transfer efficiency was highest between trophic levels 2 and 3 for both models, around 21%. Average transfer efficiency was similar in both regions; from primary producers at 15.9% on the Newfoundland and Labrador Shelf and 15.8% on the Grand Banks, and average transfer efficiency from detritus was 11.5% for the Newfoundland and Labrador Shelf and 11.8% for the Grand Banks (S5 and S6 Fig).

#### 3.2.2 Biomass and species composition

Finfish and invertebrate biomass were both slightly higher on the Newfoundland and Labrador Shelf than the Grand Banks (Fig 2, Table 1 and 2). There was higher phytoplankton biomass on the Grand Banks than the Newfoundland and Labrador Shelf, but similar amounts of zooplankton biomass. The Grand Banks had a higher number of species than the Newfoundland and Labrador Shelf, but both had similar overall fish biomass (Fig 2). The largest amount of plank-piscivorous fish biomass was on the Newfoundland and Labrador Shelf, with four times as much biomass of Arctic cod compared to the Grand Banks, where redfish dominated the plank-piscivorous fish group (Tables 1 and 2). There were higher small benthivorous fish and piscivorous fish biomasses for the Newfoundland and Labrador Shelf compared to the Grand Banks but less fish biomass for all other functional groups. The higher amount of piscivorous fish biomass for the Newfoundland and Labrador Shelf was mostly driven by Atlantic cod.

**Fig 2.**
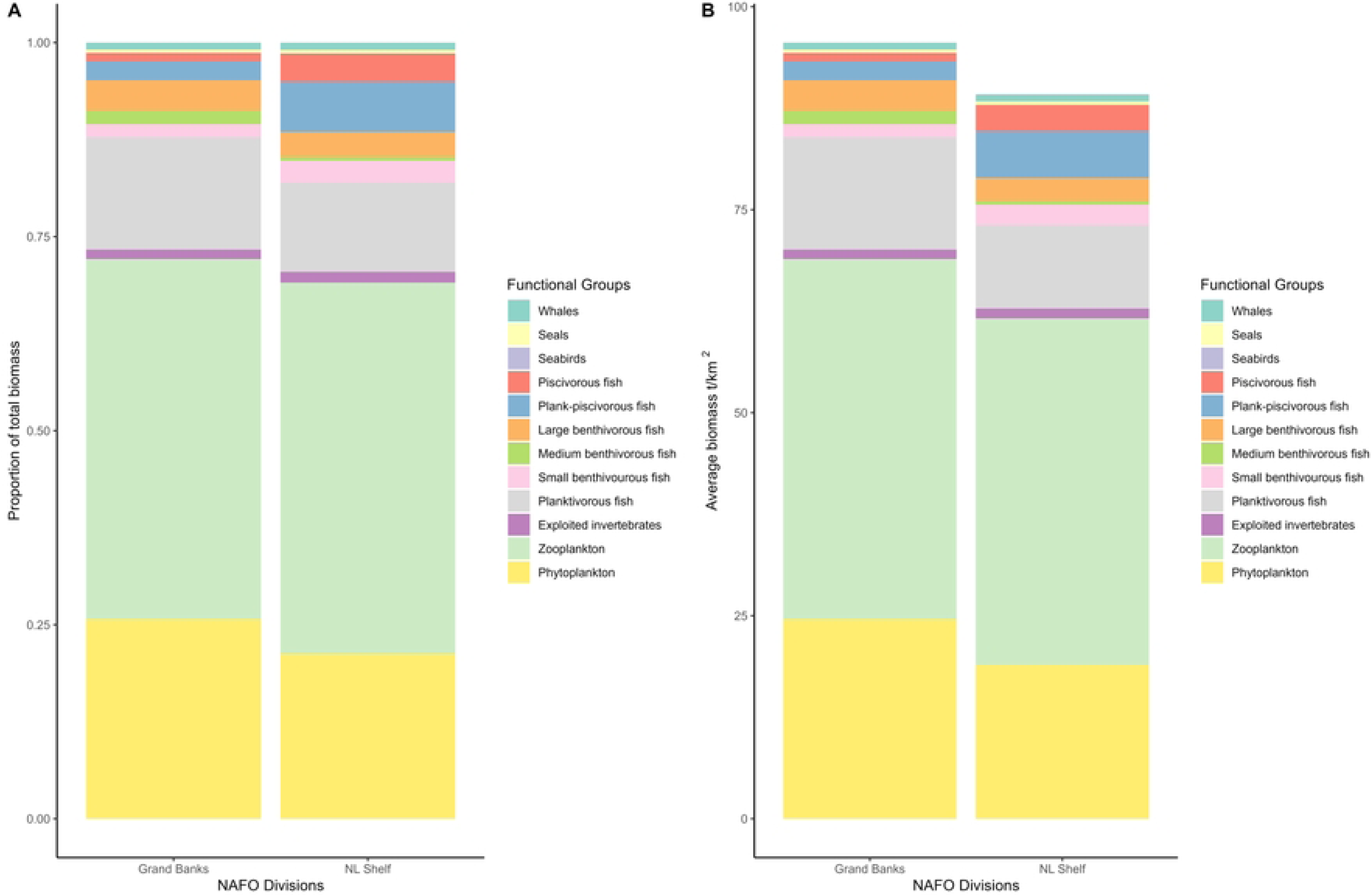
Biomass by functional group on the Newfoundland and Labrador Shelf and Grand Banks regions (2018-2020) represented by A) proportion and B) magnitude.

#### 3.2.3 Fisheries catches

In 2018-2020, there was a greater diversity and magnitude of species caught on the Grand Banks compared to the Newfoundland and Labrador Shelf (Fig 3). Proportionally, invertebrate catch contributed more to the overall percentage of catch on the Newfoundland and Labrador Shelf than the Grand Banks (46% vs 35%), but the Grand Banks still had more total invertebrate catch than the Newfoundland and Labrador Shelf (0.09 t/km^2^ vs 0.07 t/km^2^). The type of invertebrate catch was different between the two areas, with Northern shrimp dominating catch on the Newfoundland and Labrador Shelf and snow crab dominating on the Grand Banks. (S10 table). Catches on the Grand Banks for large benthivorous fish (predominantly thorny skate), medium benthivorous fish (predominantly yellowtail flounder), and plank-piscivorous fish (predominantly redfish) were almost non-existent on the Newfoundland and Labrador Shelf (Fig 3). The Newfoundland and Labrador Shelf had less capelin catch than the Grand Banks (0.011 t/km^2^ vs 0.019 t/km^2^) but the Newfoundland and Labrador Shelf had more planktivorous fish catch overall due to higher Atlantic herring (*Clupea harengus*) and Atlantic Mackerel (*Scomber scombrus*) catch. The mean trophic level of catch was 3.26 on the Newfoundland and Labrador Shelf and 3.42 on the Grand Banks (Table 3).

**Fig 3.**
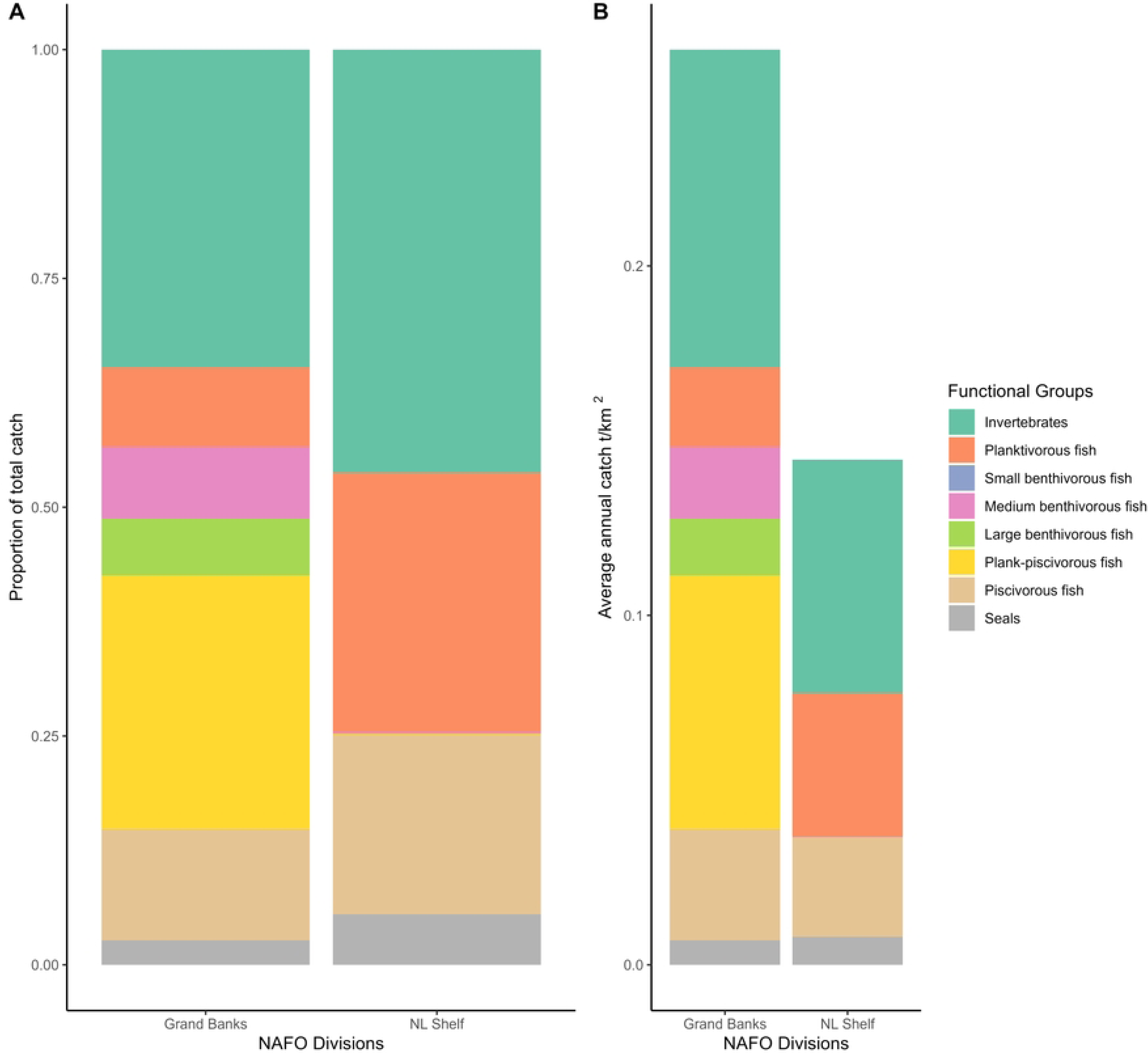
Total catches on the Newfoundland and Labrador Shelf and Grand Banks region (2018-2020) by A) proportion and B) magnitude.

**Table 3.**
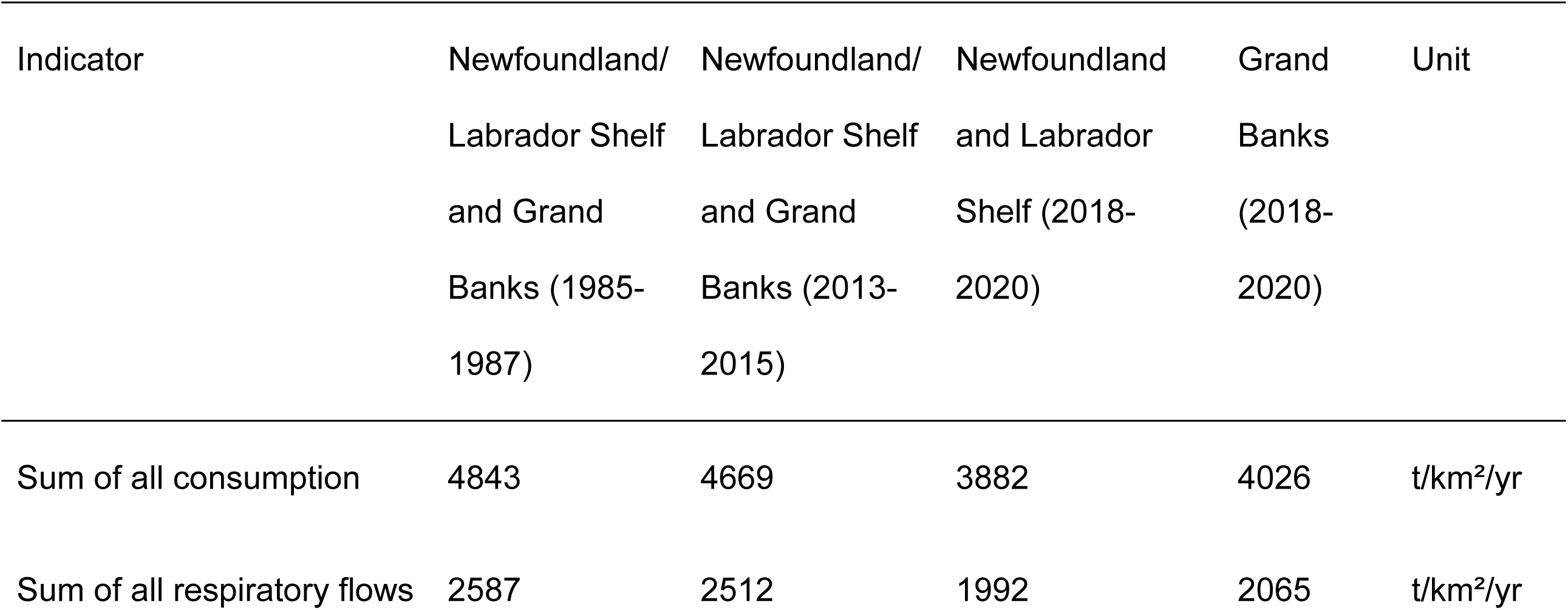
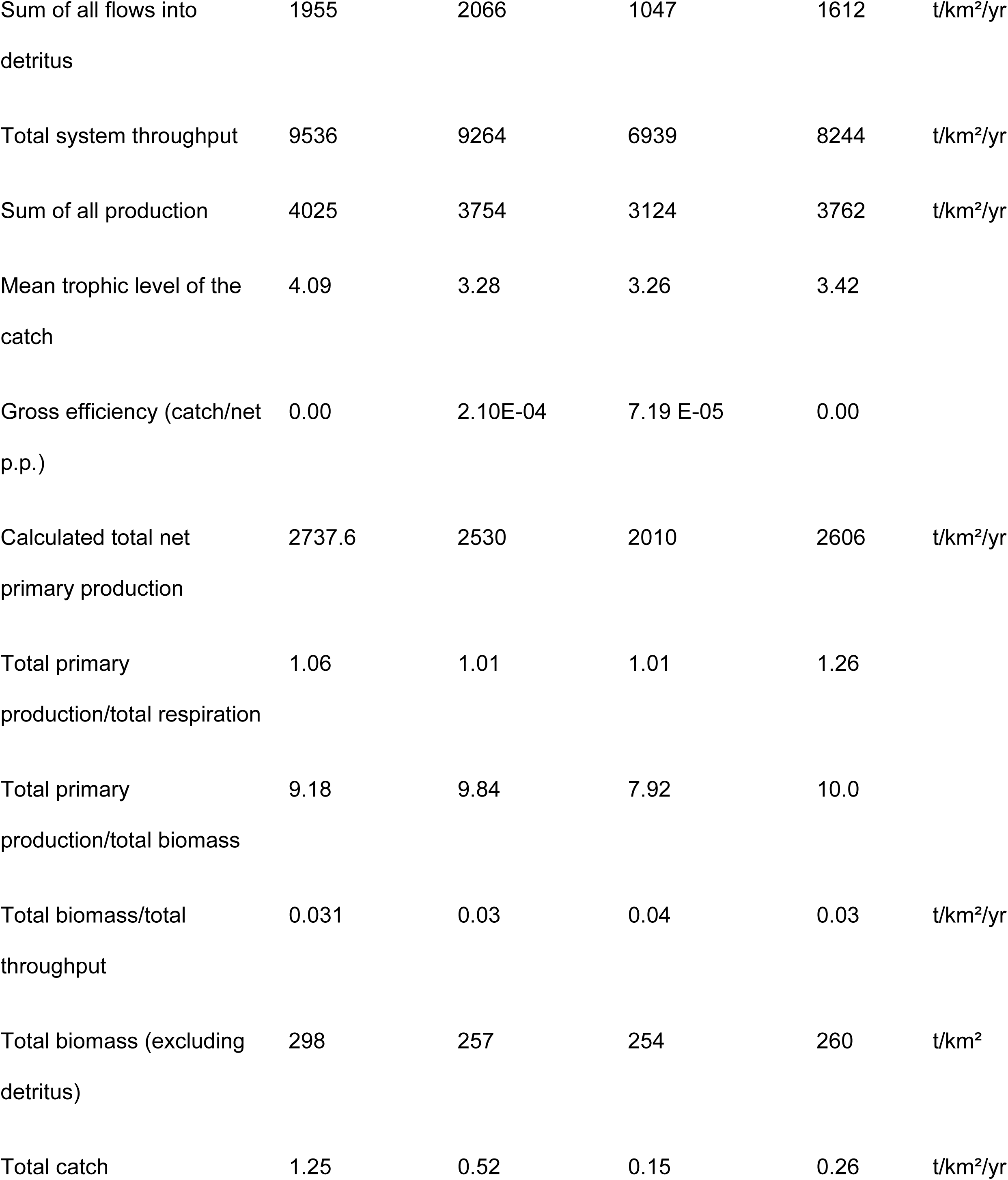
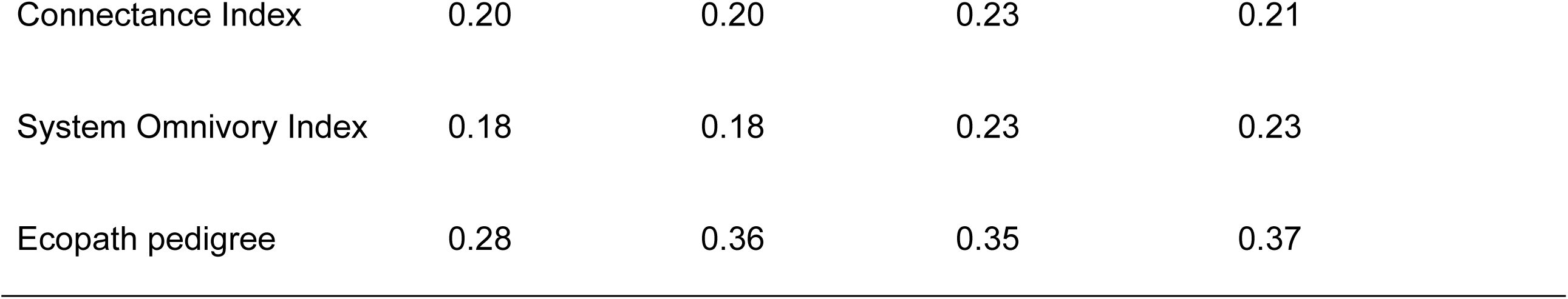
Newfoundland and Labrador Shelf and Grand Banks Ecopath model indicators for 1985-1987, 2013-2015, and 2018-2020.

| Indicator | Newfoundland/<br>Labrador Shelf<br>and Grand<br>Banks (1985-1987) | Newfoundland/<br>Labrador Shelf<br>and Grand<br>Banks (2013-2015) | Newfoundland<br>and Labrador<br>Shelf (2018-2020) | Grand<br>Banks<br>(2018-2020) | Unit |
| --- | --- | --- | --- | --- | --- |
| Sum of all consumption | 4843 | 4669 | 3882 | 4026 | t/km<sup>2</sup>/yr |
| Sum of all respiratory flows | 2587 | 2512 | 1992 | 2065 | t/km<sup>2</sup>/yr |
| Sum of all flows into detritus | 1955 | 2066 | 1047 | 1612 | t/km<sup>2</sup>/yr |
| Total system throughput | 9536 | 9264 | 6939 | 8244 | t/km<sup>2</sup>/yr |
| Sum of all production | 4025 | 3754 | 3124 | 3762 | t/km<sup>2</sup>/yr |
| Mean trophic level of the catch | 4.09 | 3.28 | 3.26 | 3.42 |  |
| Gross efficiency (catch/net p.p.) | 0.00 | 2.10E-04 | 7.19 E-05 | 0.00 |  |
| Calculated total net primary production | 2737.6 | 2530 | 2010 | 2606 | t/km<sup>2</sup>/yr |
| Total primary production/total respiration | 1.06 | 1.01 | 1.01 | 1.26 |  |
| Total primary production/total biomass | 9.18 | 9.84 | 7.92 | 10.0 |  |
| Total biomass/total throughput | 0.031 | 0.03 | 0.04 | 0.03 | t/km<sup>2</sup>/yr |
| Total biomass (excluding detritus) | 298 | 257 | 254 | 260 | t/km<sup>2</sup> |
| Total catch | 1.25 | 0.52 | 0.15 | 0.26 | t/km<sup>2</sup>/yr |

|  |  |  |  |  |
| --- | --- | --- | --- | --- |
| Connectance Index | 0.20 | 0.20 | 0.23 | 0.21 |
| System Omnivory Index | 0.18 | 0.18 | 0.23 | 0.23 |
| Ecopath pedigree | 0.28 | 0.36 | 0.35 | 0.37 |

#### 3.2.4 Mixed trophic impact

The mixed trophic impact analysis generally showed that prey groups had positive impacts on predator groups, and groups with low biomass or abundance did not have as much impact compared to more abundant species (Fig 4 and 5). Apex predators such as whales and seals generally had positive impacts on mid-trophic level species such as planktivores and benthivores that aren’t principal components of the predators diet (Fig 4 and 5). Atlantic cod >35 cm had a higher impact on the Newfoundland and Labrador Shelf compared to the Grand Banks. Groups with the greatest impact on the Newfoundland and Labrador Shelf were the positive impacts of macrozooplankton on planktivorous whales, bird, and fish species and the negative impacts of Greenland shark on seal other and whale mammal eater on other mammal groups (Fig 4). These patterns were also observed on the Grand Banks, but harp seals had greater negative impacts on their prey species and other mammal groups on the Grand Banks compared to the Newfoundland and Labrador Shelf (Fig 5).

**Fig 4.**
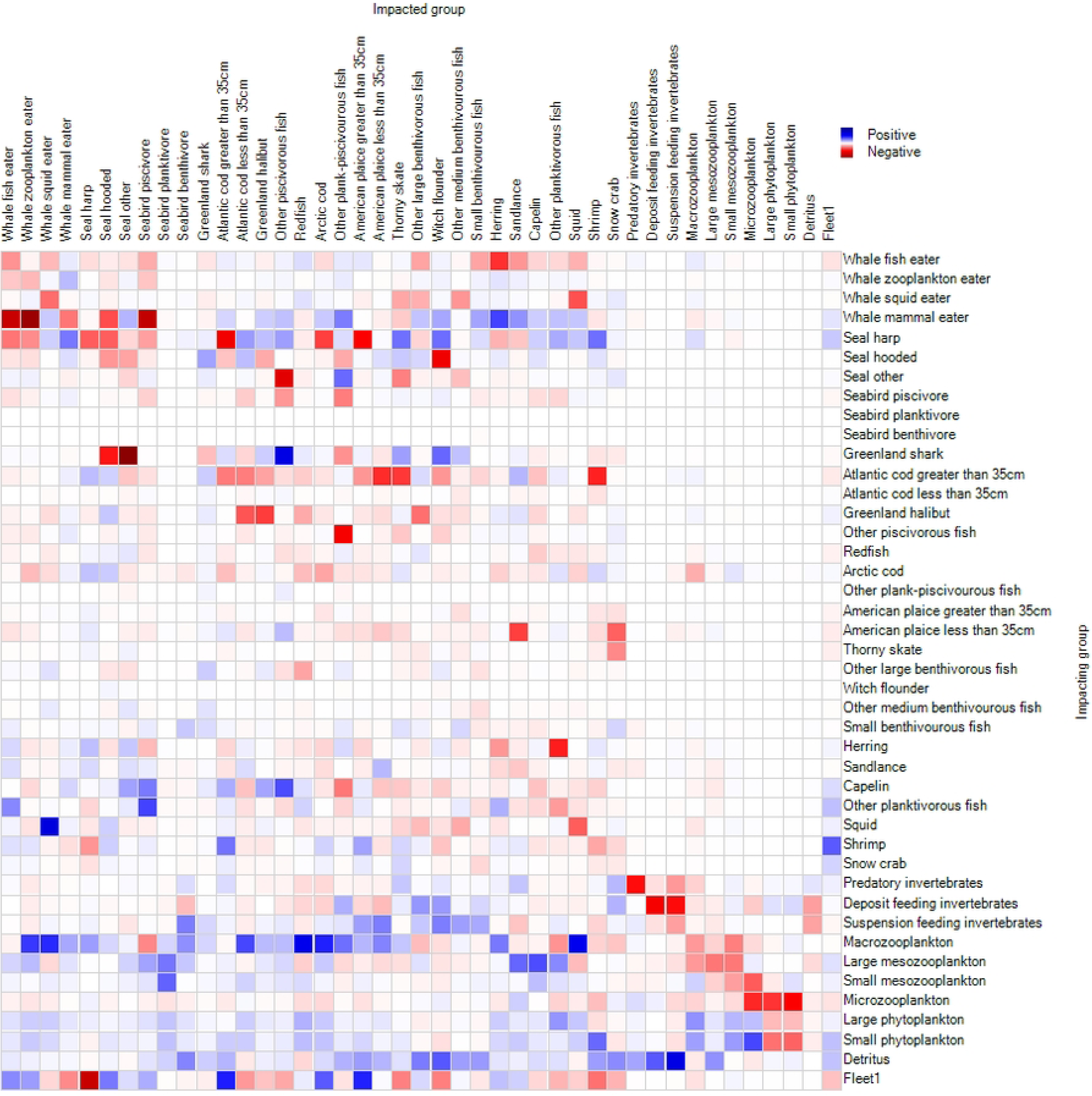
Mixed trophic impact analysis for the Newfoundland and Labrador Shelves ecosystem in 2018-2020. Functional groups (right) impacting other functional groups in the ecosystem (top). Positive impacts in blue and negative impacts in red.

**Fig 5.**
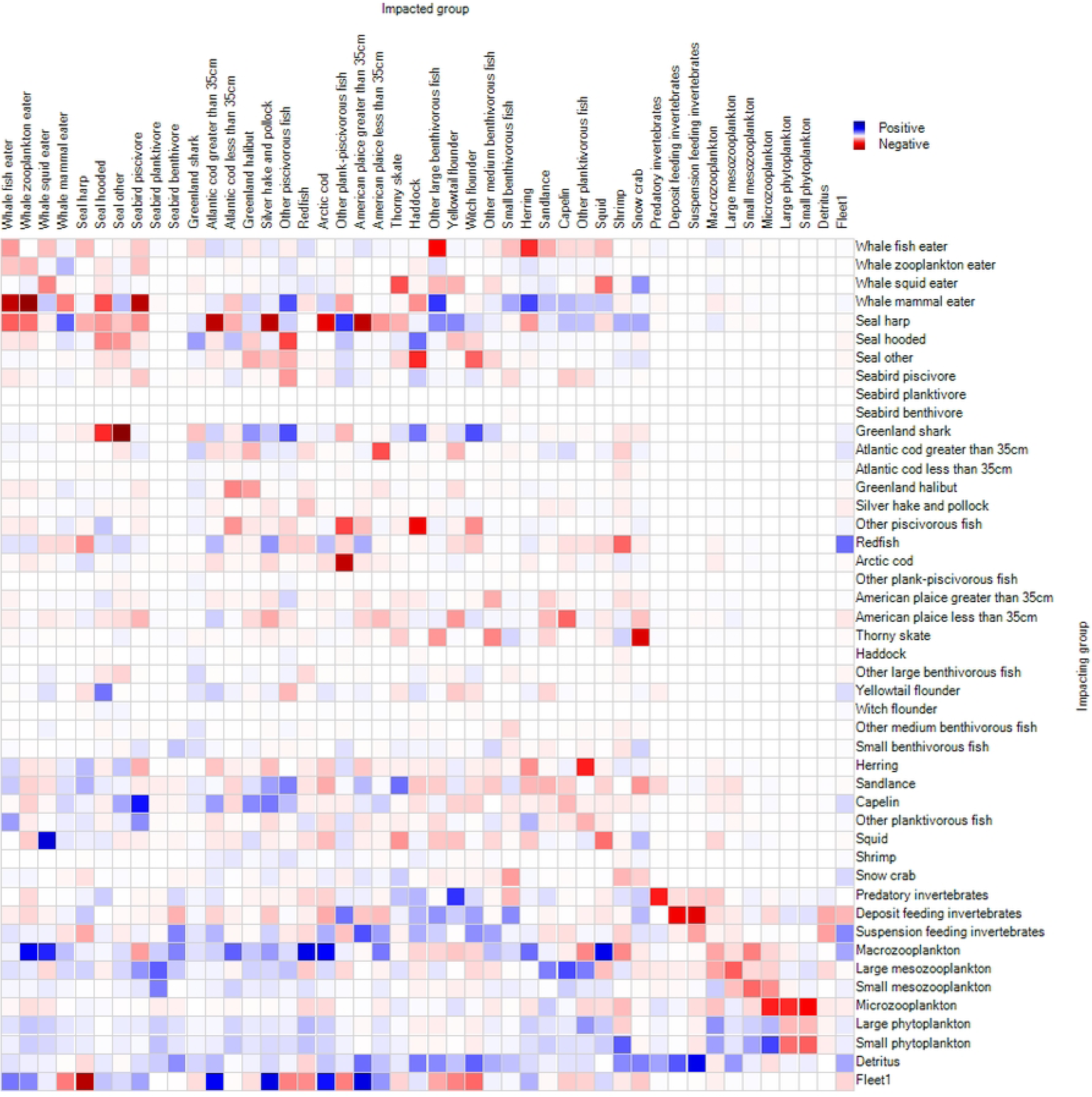
Mixed trophic impacts analysis for the Grand Banks ecosystem in 2018-2020. Functional groups (right) impacting other functional groups in the ecosystem (top). Positive impacts in blue and negative impacts in red.

In both areas, harp seals had a small positive impact on snow crab and Northern shrimp but a negative impact on Atlantic cod >35 cm (Fig 4 and 5). In both areas, capelin had a small negative impact on many medium sized benthivore fish and planktivorous fish. Capelin had a negative impact on redfish on the Grand Banks but had a small positive impact on redfish on the Newfoundland and Labrador Shelf (Fig 4 and 5). Capelin had a negative impact on all other plank-piscivorous fish groups.

### 3.3 Temporal comparisons

#### 3.3.1 Ecosystem structure and function

Total system biomass, total system catch, net primary production (NPP), net system production, and total system throughput (TST) were all higher on the Grand Banks than the Newfoundland and Labrador Shelf (Table 3). The total system omnivory index was the same for both areas, and the connectance index were similar (Table 3).

#### 3.3.2 Biomass and species composition

Total system biomass declined between 1985-1987 (298 t/km^2^) and 2013-2015 (251 t/km^2^) then increased in 2018-2020 (254 t/km^2^ for Newfoundland and Labrador Shelf and 260 t/km^2^ for Grand Banks), driven by increased secondary productivity (Fig 6B). 2013-2015 had higher invertebrate biomass than all other time periods, while 1985-1987 had the highest piscivorous and planktivorous fish biomasses (Fig 6). Finfish biomass was the highest in 1985-1987, followed by the Newfoundland and Labrador Shelf (2018–2020), the Grand Banks (2018–2020), and 2013-2015 which had the lowest. The fish community composition was different for the Newfoundland and Labrador Shelf and the Grand Banks, but with similar total biomass (Fig 6A).

**Fig 6.**
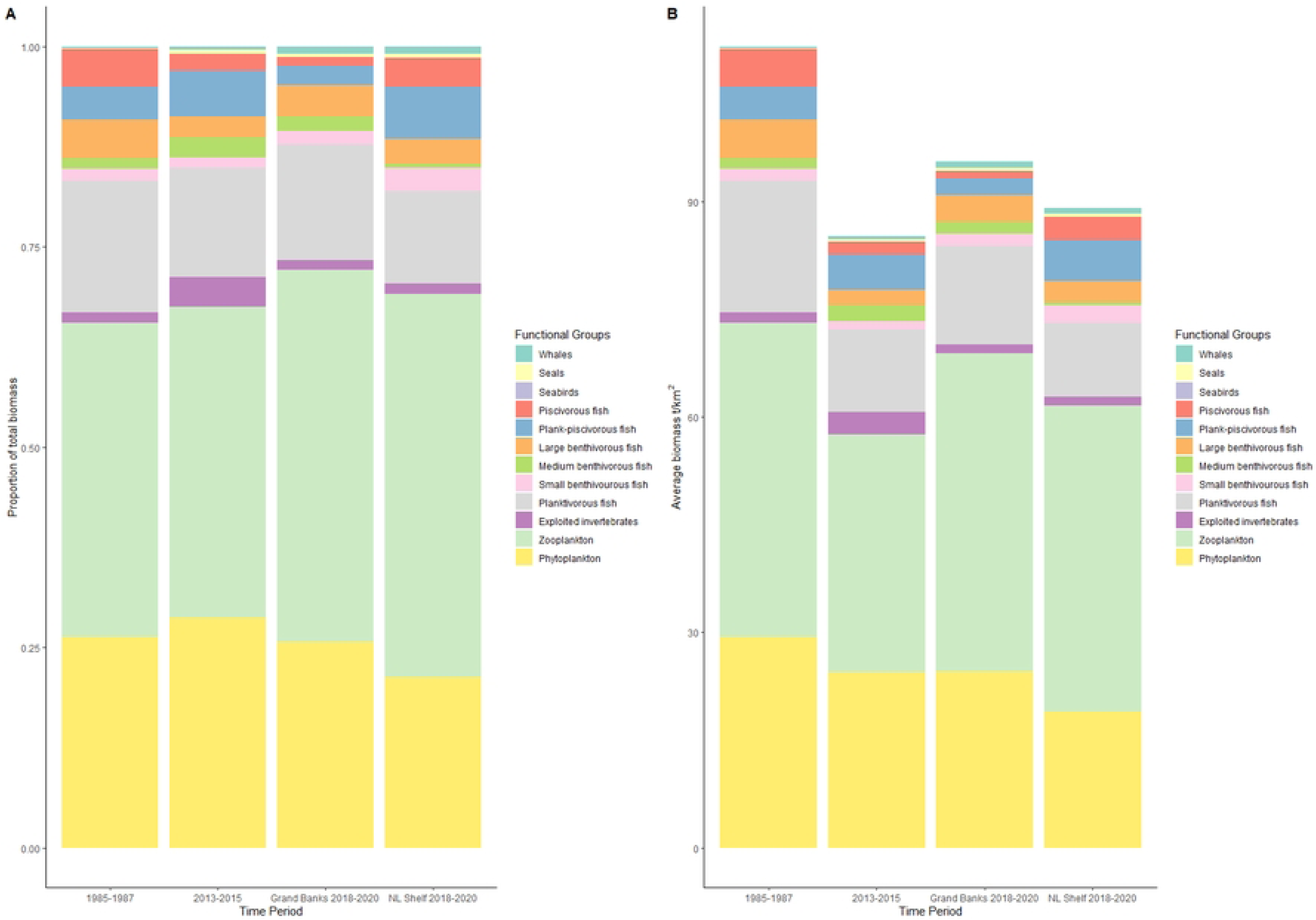
Total system biomass on the Newfoundland and Labrador Shelf and Grand Banks regions (1985-1987, 2013-2015, 2018-2020) represented by A) proportion and B) magnitude.

The highest finfish biomass was observed in 1985-1987, followed by 2018-2020 on the Newfoundland and Labrador Shelf, and similar amounts of finfish biomass in 2013-2015 and the Grand Banks in 2018-2020 (Fig 7A). The highest small benthivorous fish and piscivorous fish biomasses were observed on the Newfoundland and Labrador Shelf (2018–2020), but with lower biomass for every other fish group (Fig 7A). The increase in piscivorous fish biomass on the Newfoundland and Labrador Shelf in 2018-2020 was driven by Atlantic cod abundance, with lower Atlantic cod abundance on the Grand Banks in 2018-2020 (Tables 1 and 2). Between 1985-1987 and 2013-2015, capelin biomass declined by 64% and an additional 30% in 2018-2020 (Tables 1 and 2) (21). Sandlance, herring, and other planktivorous fish biomass remained relatively stable from 2013-2015 to 2018-2020 (Tables 1 and 2) (21). Invertebrate biomass was highest in 2013-2015, dominated by shrimp, which declined in 2018-2020 (Fig 7B). Shrimp biomass on the Newfoundland and Labrador Shelf and Grand Banks in 2018-2020 was less than in 1985-1987 (Fig 7B). Snow crab and squid biomasses on the Grand Banks in 2018-2020 were the highest of all periods, while shrimp biomass on the Grand Banks was the lowest of all periods (Fig 7B). Total plankton biomass declined between 1985-1987 and 2013-2015 and increased in both the Newfoundland and Labrador Shelf and the Grand Banks for 2018-2020 (Fig 7C). This was largely due to increases in macrozooplankton and large mesozooplankton, comprised of non-pandalus shrimp, gelatinous zooplankton, and most importantly, large copepod species such as *Calanus finmarchicus*. Small mesozooplankton and microzooplankton biomasses were constant across time periods. Large and small phytoplankton biomasses generally decreased through the time periods, except for the Grand Banks in 2018-2020, with increased small phytoplankton biomass (Fig 7C).

**Fig 7.**
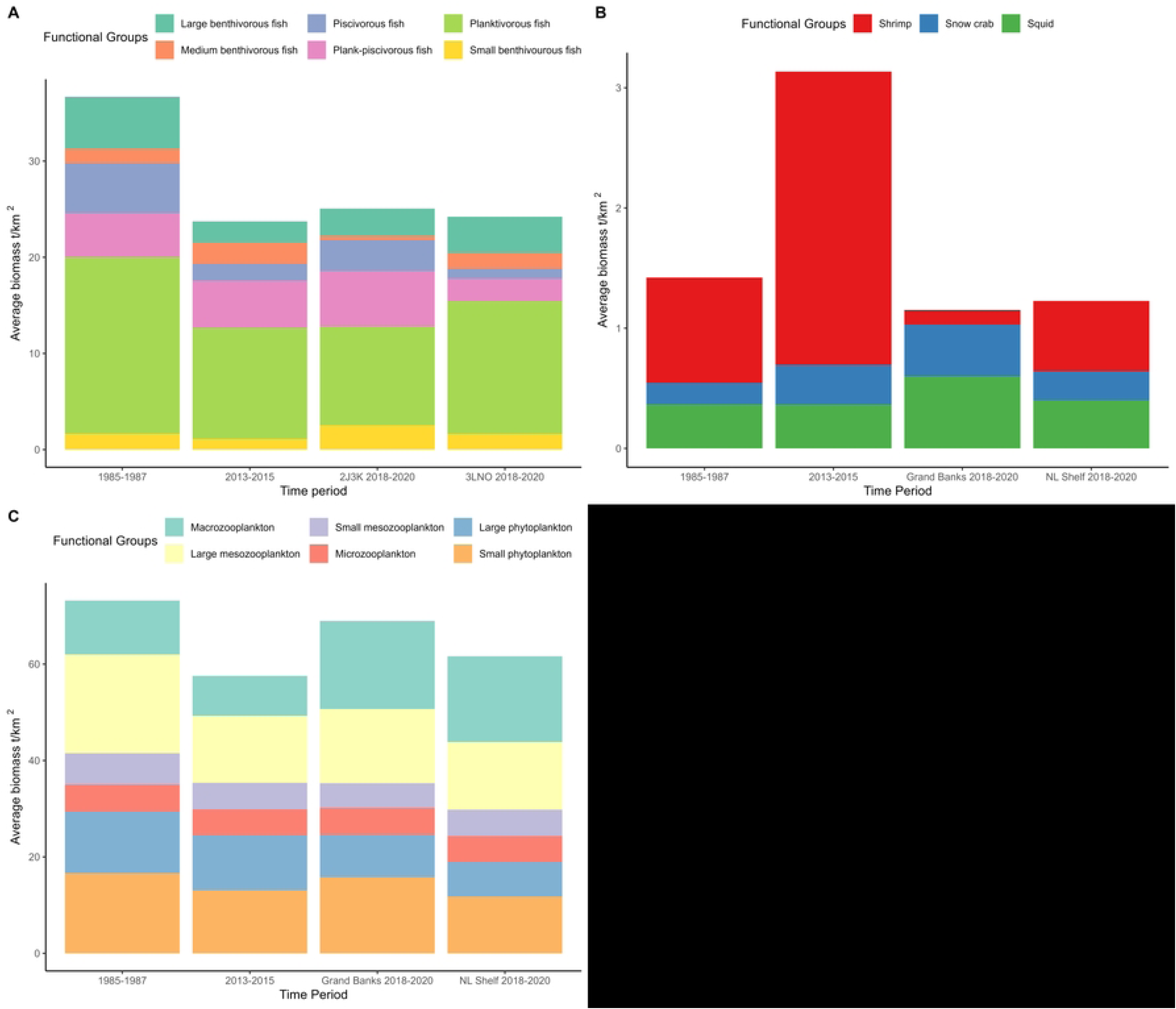
Biomass (t/km²) for the 1985-1987 and 2013-2015 Newfoundland Shelf/Grand Banks combined models, and the 2018-2020 Newfoundland Shelf and 2018-2020 Grand Banks model for A) Finfish B) Invertebrates and C) Plankton.

#### 3.3.3 Fisheries catches and mortality

There was a large drop in total catch and a change in the proportion of species being caught through time periods (Fig 8). For both the Newfoundland and Labrador Shelf and the Grand Banks, between 1985-1987 and 2013-2015, piscivorous fish (predominantly cod) catches declined and invertebrate (predominantly northern shrimp and snow crab) catches increased (Fig 8). On the Newfoundland and Labrador Shelf, catch declined by approximately 0.2 t/km^2^ between each time period. On the Grand Banks, the initial decline between 1985-1987 and 2013-2015 was larger (approximately 0.4 t/km^2^) than the decline between 2013-2015 and 2018-2020 (approximately 0.07 t/km^2^). This trend in overall catch decline was reflected in declining fishing mortality rates for all three time periods (Fig 9). In 2018-2020, no functional group experienced a higher fishing mortality rate than natural mortality, whereas 1985-1987 cod and 2013-2015 snow crab had higher fishing mortality than natural mortality (Fig 9). In all four models, squid had the highest natural mortality rate out of the commercially harvested species (Fig 9). On average, natural mortality rates increased from 1985-1987 to 2013-2015, and then remained stable from 2013-2015 to 2018-2020. The Newfoundland and Labrador Shelf and Grand Banks both had similar levels of natural mortality across functional groups, with more variability appearing in some groundfish species such as Greenland halibut and American plaice (Fig 9, S11 and S12 Table).

**Fig 8.**
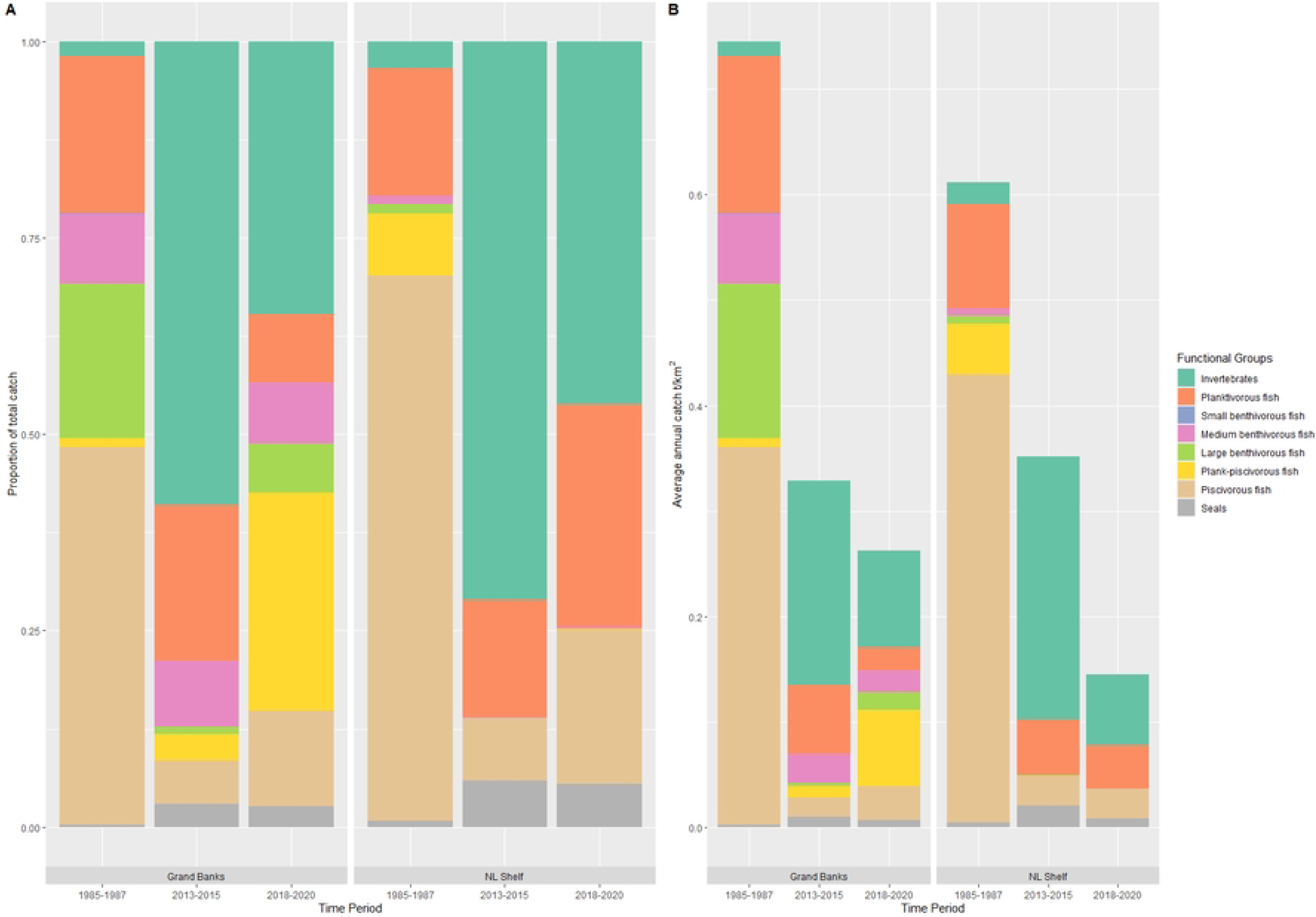
Catches on the Newfoundland and Labrador Shelf and Grand Banks (1985-1987, 2013-2015, and 2018-2020) represented by A) proportion and B) magnitude.

**Fig 9.**
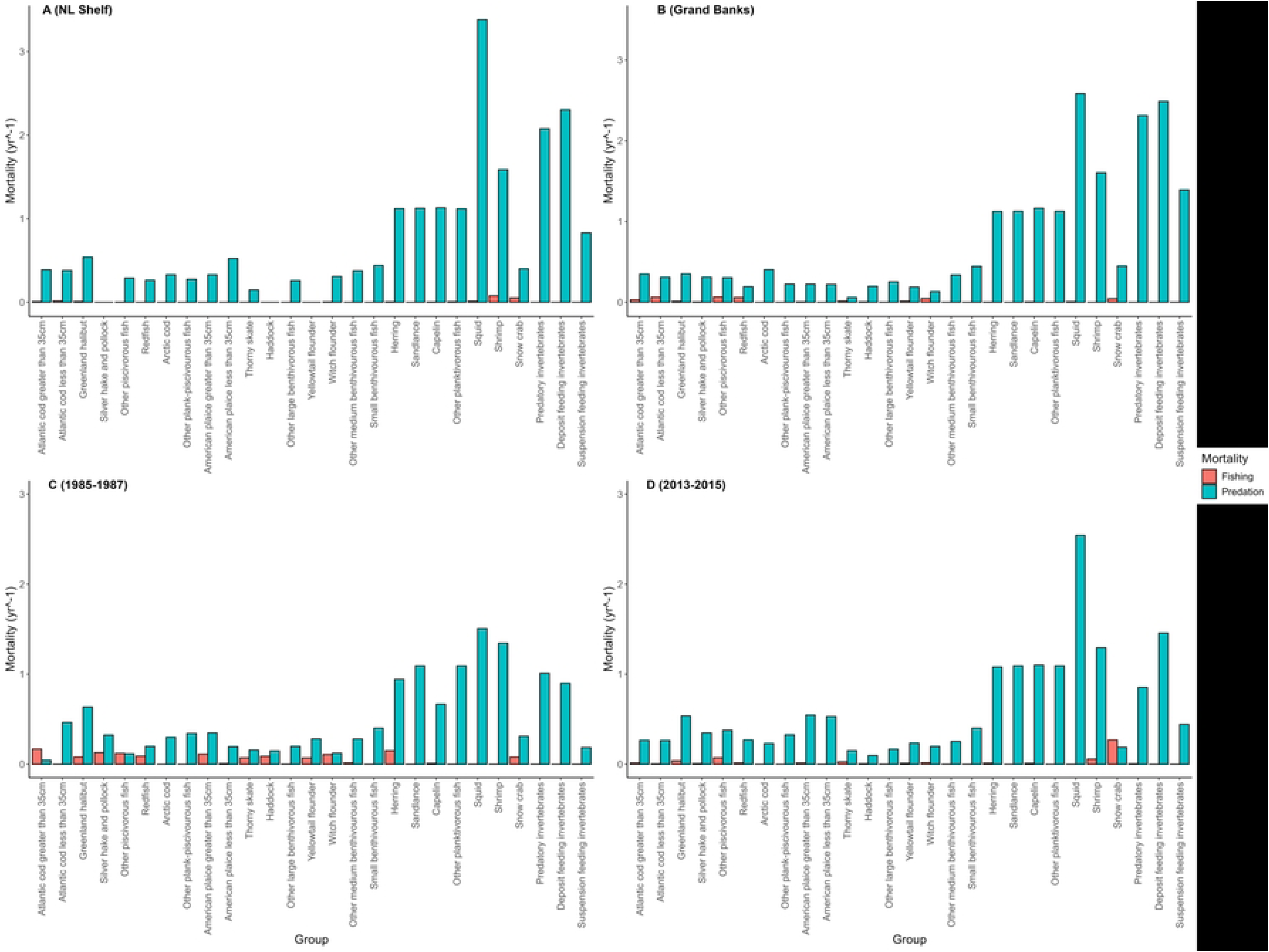
Fishing and predation mortalities for commercial species in A) Newfoundland and Labrador Shelf 2018-2020 B) Grand Banks in 2018-2020 C) Newfoundland and Labrador Shelf and Grand banks 1985-1987 and D) Newfoundland Shelf and Grand Banks 2013-2015.

## Discussion

### 4.1 Summary

Ecosystem structure and function varied between the Newfoundland and Labrador Shelf and the Grand Banks in 2018-2020. The Grand Banks had higher species richness and species diversity than the Newfoundland and Labrador Shelf. Despite having lower finfish and invertebrate biomass, higher primary and secondary productivity contributed to the Grand Banks having higher total system biomass than the Newfoundland and Labrador Shelf. Species such as Arctic cod, Atlantic cod, and Northern shrimp were more abundant on the Newfoundland and Labrador Shelf, while sandlance, redfish, and snow crab were more abundant on the Grand Banks. Overall catch rates were nearly double on the Grand Banks compared to the Newfoundland and Labrador Shelf, with greater catches on the Grand Banks for almost every functional group except Northern shrimp, Greenland Halibut, Atlantic herring, and other planktivorous fish.

Ecosystem structure and function also varied across time. Overall system biomass increased from 2013-2015 to 2018-2020, but was lower than the 1985-1987 period. The only group this pattern did not hold for was exploited invertebrates, which peaked in 2013-2015. Species abundances varied across time periods. The last two time periods are five years apart, indicating that the region is undergoing a period of rapid change. Catch levels declined from 1985-1987 to 2013-2015, and again in 2018-2020. The high invertebrate catch rates of 2013-2015 declined in 2018-2020, while planktivorous and piscivorous fish catches for Atlantic herring, capelin and Atlantic cod increased.

### 4.2 The role of forage species

Forage fish play a crucial role in food webs and ecosystem dynamics globally, by providing a link between plankton groups and large predators (79). In some cases, this important role can create a ‘wasp-waist’ ecosystem, where the link between low and high trophic levels is limited to one or a few key species (80,81). In the Newfoundland and Labrador ecosystem, capelin is a forage fish with a large influence on high trophic levels. Seasonal humpback whale migrations are linked to capelin, with whales returning to the same foraging sites each year (82). Harp seal and seabird fecundity rates on the Grand Banks improve with high capelin abundance (83,84). Atlantic cod mortality rates and overall productivity are highly linked with capelin on the Newfoundland and Labrador Shelf (13). Although the cause of the capelin collapse remains unclear, their population dynamics seem to be driven primarily by bottom-up forces, such as the influence of sea-ice on the spring bloom (15,85).

With reduced capelin abundance in the two recent time periods, both study areas had an alternate key forage fish that was abundant in biomass, unexploited by commercial fisheries, and was prominent in predator diets – Arctic cod on the Newfoundland and Labrador Shelf and sandlance on the Grand Banks (86). Squid was also an important alternative prey group in the models with a spike in its abundance in the late 2010s (87). Other shifts in predator diets have been observed following the decline in capelin, such as Greenland halibut shifting toward shrimp (88), gulls shifting toward a more generalist diet including other birds’ eggs (89,90), and Atlantic cod to shrimp and crab (91). Humpback whale diets remain dominated by capelin despite their reduced availability (82).

### 4.3 Highly dynamic ecosystems

Developing ecosystem models during periods of relative ecosystem stability can help to better understand ecosystem dynamics. The boom of capelin biomass observed in 2013-2015 declined in 2018-2020, and despite this Atlantic cod biomass increased in the same time frame. A similar situation occurred in the Barents Sea, which historically had Atlantic cod as a dominant predator and the main fished species, with collapses or declines in groundfish and forage fish in past decades (92–94). The cod population in the Barents Sea has recovered from decline despite frequent collapses in capelin populations (94). The lack of capelin and cod recovery in Newfoundland and Labrador has been attributed to greater magnitude of decline and more intensive historical fishing pressure (92,95).

The Gulf of Alaska is another highly dynamic system that is heavily influenced by environmental drivers (10). In 2014-2016, the Gulf of Alaska experienced a prolonged period of high sea surface temperature, which greatly reduced system productivity and increased metabolic demands on many key species, including a 70% decline in Pacific cod stocks over three years (96). EBFM management strategies included a ban on forage fish catch, focusing on system level groundfish yield, as opposed to single species, and inclusion of stakeholders in industry and science in the decision-making process. These EBFM practices mitigated biological and socio-economic losses, and provide a template for adapting fisheries to climate change (97). A similar management strategy to ban commercial harvest of capelin could be an option to explore for its ability to promote the recovery of capelin and cod in Newfoundland and Labrador.

### 4.4 Model limitations and assumptions

#### 4.4.1 Natural variability

Seasonal, interannual, or ontogenetic variability in consumption and production rates can introduce error into models (98). EwE models provide static snapshots of annual averaged rates that may not reflect the peaks and valleys of parameters over time. For example, phytoplankton blooms occur in the spring and fall with high amounts of biomass, and less biomass in the winter and summer. Seasonal migrations can also introduce variability. For example, harp seals migrate in and out of both model domains seasonally as a part of two herds, one into the Gulf of St. Lawrence, and the other around the Newfoundland and Labrador Shelf (48). Harp seals have an approximate residence time of 228 days, or ∼60% of the year in the study area and ∼40% spent in the Arctic, but the timing can change with sea ice conditions (49). There was no estimate available to split the total harp seal biomass between the Newfoundland and Labrador Shelf and Grand Banks. It was assumed that there was an equal division of harp seal biomass between both areas, despite the understanding that harp seals seem to occur more frequently in the Newfoundland and Labrador shelf as opposed to the Grand Banks (49–51).

#### 4.4.2 Observation error

Observation error arises from imperfect methods of observation when measuring or sampling (98). For example, phytoplankton size classes derived from remote sensing may under-represent smaller phytoplankton, and do not differentiate between ‘large’ and ‘small’ cell sizes with 100% accuracy (99). The stratified random sampling approach by RV trawl surveys assumes individuals are drawn randomly and equally from the population, however small pelagic and semi-demersal species tend to be underestimated in these samples as these species are not reliably caught in a bottom trawl. Sandlance and Arctic cod are known to be under-represented in RV survey catches and whose biomass had to either be estimated by Ecopath or scaled based on acoustic surveys. Due to high dietary demand from predators, the 2018-2019 model may have unrealistically high estimates of other forage fish biomass such as Atlantic herring in order to achieve mass balance.

#### 4.4.3 Modelling framework

The models developed in this study represent two hypotheses of many potential ecosystems. The complexity of ecosystem models can introduce variance when parameters are estimated that lack robust empirical data (100). Our rationale for removing the bacterial loop in the 2018-2020 models stemmed from this concern. Uncertainty in input parameters highlights knowledge gaps for some important species. Some data-poor groups used biomass estimates from the 2013-2015 model, such as hooded seal, other seals, suspension feeding/deposit feeding/predatory invertebrates, and seabird functional groups (21). These assumptions may bias the 2018-2020 models to appear more similar to the 2013-2015 model than the ecosystem is in reality.

### 4.5 Future Applications

The recovery of overexploited fish populations requires that catch rates remain low or at zero, especially for forage fish (79,81). While a reduction in fishing pressure can produce large shifts in ecosystem structure (101,102), environmental conditions can confound this relationship (103). Variability in sea surface and bottom temperatures, among other environmental variables could be influencing primary production and energy transfer to high trophic levels, impacting species recovery (104). In the Barents Sea, a multi-model approach to fisheries where complimentary ecosystem models explored different aspects of trophic interactions between cod, capelin, polar cod, and copepods under varying amounts of fishing pressure found that indirect food-web effects might be just as important predator-prey interactions (105). A similar approach could be applied for the Grand Banks and Newfoundland and Labrador Shelf ecosystems to explore scenarios under various primary production, fishing, diet configurations, and predator states. These scenarios could shed light on what conditions would be necessary to facilitate a forage fish recovery in the system, and if that would result in an overall system biomass recovery. Such findings would be invaluable to management strategy evaluations and EBFM applications seeking to improve fisheries management in highly dynamic ecosystems (18,106). A deeper understanding of the factors that regulate the ecosystem will be critical to successfully manage fished stocks, in highly dynamic ecosystems under climate change.

## 5. Conclusions

The value of synthesizing information to describe ecosystem structure and function lies in the approach to explore ecological hypotheses. Ecosystem structure and function of the Newfoundland and Labrador Shelf and the Grand Banks varied spatially, and temporally, with shifts from the 1980s to the late 2010s. Fisheries catches have declined since 1985-1987 and are lower on the Newfoundland and Labrador Shelf compared to the Grand Banks. Changes in ecosystem structure and function and fisheries target species observed in the short time from 2013-2015 to 2018-2020 indicates that the system has recently undergone a period of rapid change. While a modest increase in groundfish may not be indicative of a full system recovery, increases in zooplankton biomass in combination with a decline in invertebrate biomass could set the stage for a further shift away from invertebrate dominance, and toward increasing groundfish and forage fish stocks in the next decade.

## Acknowledgements

We thank Alida Bundy and Maxime Geoffroy for their constructive comments on the MSc thesis that formed the basis for this study.

We also thank Mariano Koen-Alonso, Carina Gjerdrum, and Shelley Lang for providing data and interpretation used to inform the model.

## Supporting Information

**S1 Table. List of parameter estimates and relevant literature for the Newfoundland and Labrador Shelf** (**2018–2020**) **model.**

**S2 Table. List of parameter estimates and relevant literature for the Grand Banks** (**2018–2020**) **model.**

**S3 Table. Diet matrix for the Newfoundland and Labrador Shelf model** (**2018–2020**).

**S4 Table. Diet matrix for the Grand Banks model** (**2018–2020**).

**S5 Table. Data pedigree for the Newfoundland and Labrador Shelf model** (**2018–2020**). Higher numbers and darker shading indicate higher uncertainty.

**S6 Table. Data pedigree for the Grand Banks model** (**2018–2020**). Higher numbers and darker shading indicate higher uncertainty.

**S7 Table. Unbalanced Newfoundland and Labrador Shelf model** (**2018–2020**). Values in blue are estimated by Ecopath. Bolded Values were changed in the balancing process.

**S8 Table. Unbalanced Grand Banks model** (**2018–2020**). Values in blue are estimated by Ecopath. Bolded values were changed in the balancing process.

**S9 Table. Common and latin names of species included in each functional group for the Newfoundland and Labrador Shelf and Grand Banks** (**2018–2020**) **models.**

**S10 Table. Total catch (t/km^2^) of each functional group for the Newfoundland and Labrador Shelf and Grand Banks** (**2018–2020**) **models.**

**S11 Table. Predation mortality rates across all functional groups for the Newfoundland and Labrador Shelf model** (**2018–2020**).

**S12 Table. Predation mortality rates across all functional groups for the Grand Banks model** (**2018–2020**).

**S1 Fig. Change in parameter estimates (%; B = biomass, P/B = production: biomass, Q/B = consumption: biomass) between the Newfoundland and Labrador Shelf 2018-2020 unbalanced and balanced models.** The darkest blue squares include values that are greater than -200%.

**S2 Fig. Percent change in parameters (%; *B =* biomass, *P/*B = production: biomass, Q/B = consumption: biomass) between the Grand Banks 2018-2020 unbalanced and balanced models. The darkest blue squares include values that are greater than -200%.**

**S3 Fig. Percent change in the 2018-2020 Newfoundland and Labrador Shelf model diet matrix between the balanced and unbalanced model.**

**S4 Fig. Percent change in the 2018-2020 Grand Banks model diet matrix between the balanced and unbalanced model.**

**S5 Fig. Lindeman spine of trophic flows on the Newfoundland and Labrador Shelf ecosystem in 2018-2020. Roman numerals represent trophic levels (TL). Flows to detritus are recycled through the detritus and primary production (D+P) comportments as trophic level I.**

**S6 Fig. Lindeman spine of trophic flows on the Grand Banks ecosystem in 2018-2020. Roman numerals represent trophic levels (TL). Flows to detritus are recycled through the detritus and primary production (D+P) comportments as trophic level I.**

